# Beyond body maps: information content of specific body parts is distributed across the somatosensory Homunculus

**DOI:** 10.1101/2021.08.23.457376

**Authors:** Dollyane Muret, Victoria Root, Paulina Kieliba, Danielle Clode, Tamar R. Makin

## Abstract

The somatosensory homunculus in primary somatosensory cortex (S1) is topographically organised, with relatively high selectivity to each body part in its primary area. This dominant feature may eclipse other organising principles in S1. Recent multivariate methodologies allow us to identify representational features beyond selectivity, e.g., information content, providing new opportunities to characterise the homunculus. Using Representational Similarity Analysis, we asked whether body part information content can be identified in S1 beyond the primary area of a given body part. Representational dissimilarities in fMRI activity patterns were compared between different body parts (face, hand and feet) and subparts (e.g., fingers), and between different actions performed with the same body part. Throughout the S1 homunculus, we identified significant dissimilarities between non-primary body parts (e.g., between the hand and the lips in the foot area). We also observed significant dissimilarities between body subparts in distant non-primary areas (e.g., different face parts in the foot area). Finally, we could significantly dissociate between two movements performed by one body part (e.g., the hand) well beyond its primary area (e.g., in the foot and face areas), even when focusing the analysis along the most topographically organised sub-region of S1, Brodmann area 3b. Altogether, our results demonstrate that body part and action-related information content is more distributed across S1 homunculus than previously considered. While this finding does not revoke the general topographic organising principle of S1, it reveals yet unexplored underlying information contents that could be harnessed for rehabilitation, as well as novel brain-machine interfaces.

## Introduction

Contrary to its motor counterpart, the primary somatosensory cortex (hereafter S1) is considered as highly topographically organised, with relatively high levels of selectivity within each body part‘s representation (Schieber, 2001; Cunningham *et al*., 2013; Huber *et al*., 2020). This perspective over S1 organisation arises from a long-lasting mapping tradition, initiated in the 19^th^ century (Fritsch and Hitzig, 1870; Ferrier, 1873; Penfield and Boldrey, 1937) and continued since then in electrophysiology (Merzenich *et al*., 1978; Kaas *et al*., 1979; Baldwin, Cooke and Krubitzer, 2017), cortical stimulation (Roux, Djidjeli and Durand, 2018; Sun *et al*., 2021) and neuroimaging studies (Nakamura *et al*., 1998; Germann *et al*., 2020; Saadon-Grosman, Arzy and Loewenstein, 2020; Willoughby, Thoenes and Bolding, 2021). This conventional mapping approach assigns brain function to a given cortical area by selecting the most responsive body part for a set of neurons or voxels, in a winner-takes-all manner. While this approach has been hugely beneficial, e.g. for understanding (Kaas *et al*., 1979) and restoring (Bensmaia and Miller, 2014; Flesher *et al*., 2016) brain function, it may also eclipse the presence of other organising principles which may also bare relevance for S1 function. In particular, the underlying (weaker) inputs that tend to be neglected in mapping approaches, may also provide functional contributions, even if secondary to the dominant input.

Recent methodological advancement (e.g., multivariate pattern analysis) allow to identify representational features beyond selectivity at the single unit (or voxel) level, and thus provide new opportunities to characterise the homunculus. Using complementary approaches, a recent study using direct neural recording in tetraplegic patients revealed the presence of latent activity evoked by movements from the entire body in the hand area of the primary motor cortex (hereafter M1) (Willett *et al*., 2020). These results were reported in a pathological context, but they are consistent with the notion that other organising principles beyond somatotopy, e.g. representation of ethologically relevant actions (Graziano and Aflalo, 2007), may underlie the general organisation of body related motor maps. While this work provided a new perspective over the intrinsic organisation of M1, the organising principles of S1 remain so far relatively unquestioned. Yet we know that M1 and S1 are tightly coupled, both anatomically and functionally (Matyas *et al*., 2010; Catani *et al*., 2012; Kumar, Manning and Ostry, 2019). Interestingly, the M1 and S1 hand areas were found to share similar representational features (Ejaz, Hamada and Diedrichsen, 2015; Wesselink *et al*., 2019), which are in both cases better explained by inter-finger co-use in daily life than by topographic organisation (Ejaz, Hamada and Diedrichsen, 2015). Thus, despite their physiological differences, S1 and M1 may share more functional organisation than previously thought, especially in terms of information content. This raises the question of whether S1 could also contain additional representational patterns underlying its topographic organisation.

Recent evidence indeed points towards a more complex organisation of S1, beyond its topographic organisation. For example, an imaging study of the negative BOLD responses in humans revealed the presence of an underlying inverted homunculus (Tal, Geva and Amedi, 2017), similar to M1 (Zeharia *et al*., 2012), suggesting a distribution of activity patterns across the Homunculus. In addition, recent reports in rodents show that the information content arising from different tactile inputs provided to a digit, could be decoded even from a non-adjacent digit representation (Enander and Jörntell, 2019). This and other recent evidence (Tommerdahl, Favorov and Whitsel, 2010; Thakur, Fitzgerald and Hsiao, 2012; Schellekens *et al*., 2020; Wesselink *et al*., 2020) stress the need to investigate the distribution of representational information content throughout S1 homunculus.

Here we investigate whether information content can be identified in S1 beyond the primary area of a given body part, as defined by conventional mapping criteria. We asked healthy participants to perform a series of sensorimotor paradigms in the scanner: i) individual fingers movements (hereafter finger task), ii) movements of specific facial parts (hereafter face task), or iii) two different actions (i.e., squeeze or push an object) with each of three body parts (i.e., lips, hand, feet; hereafter body task). Using conventional univariate analyses on an independent dataset, we first defined individual S1 regions of interest showing high selectivity to face, hand and foot movements. We then used Representational Similarity Analysis (RSA) to index information content by quantifying multivoxel representational dissimilarities between actions and body parts. Cross-validated Mahalanobis distances provide a quantification of these dissimilarities, where distances that are greater from zero reflect significant information content (note that we deliberately avoid the term ‘representational content’ since it could imply functional relevance). We found task-relevant information content was distributed across S1, demonstrating an intrinsic organisation to S1 beyond somatotopy.

## Results

Our main approach was to identify distant and highly selective regions of the S1 homunculus at the individual participant level, to assess their univariate activity levels and multivariate information content. Towards this end, within an S1 anatomical landmark (black contours in Fig. 1A) we defined three regions of interest (hereafter ROIs) for each participant in each hemisphere, based on the 50 most selective voxels from an independent body localiser of the foot, hand and face (see Fig. 1A for consistency maps across participants).

**Figure 1.**
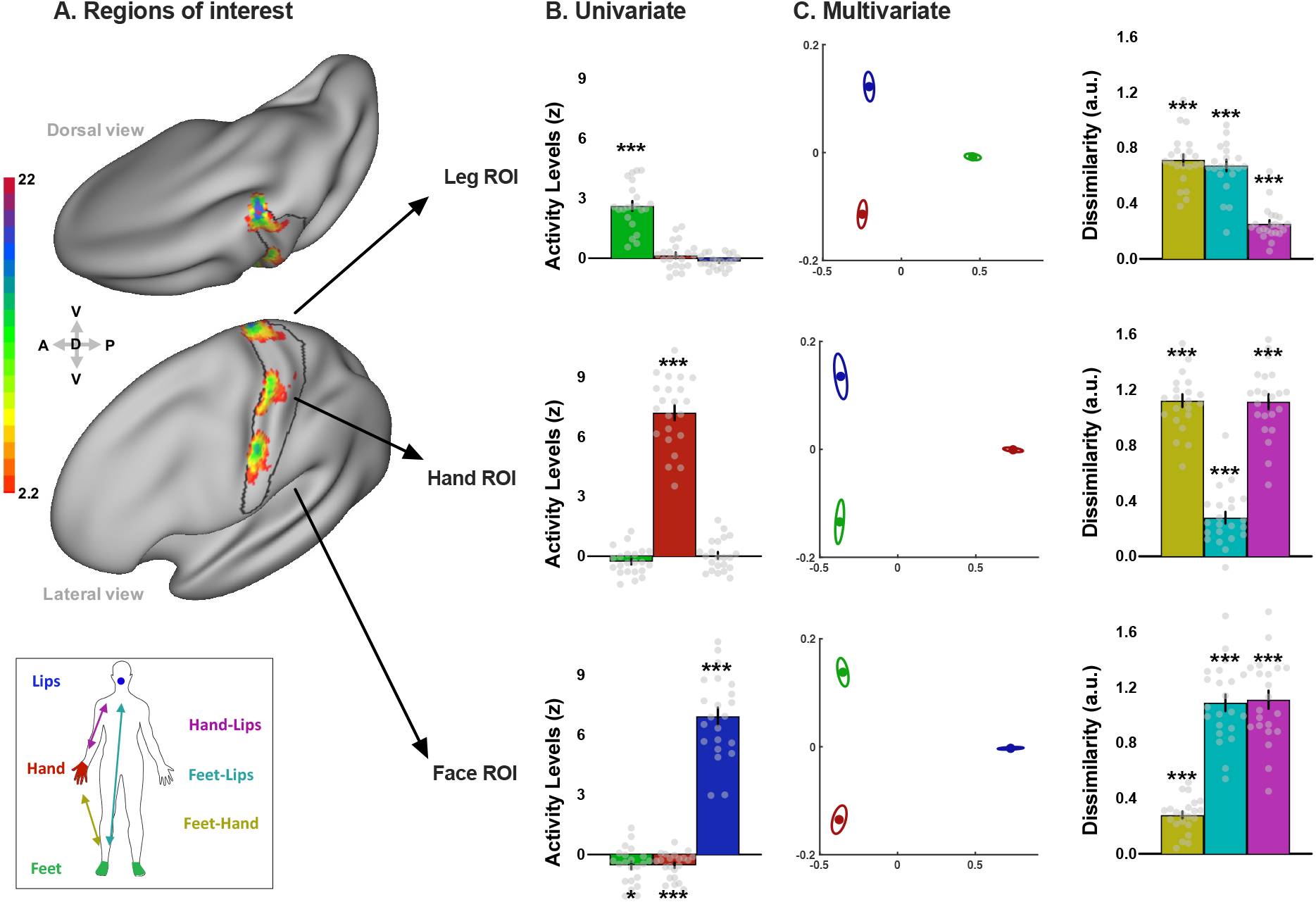
Regions of interest, selectivity, and multivariate information content related to specific body parts across S1 Homunculus. **A)** Consistency maps across participants of the S1 regions of interest (ROIs) for the body task (n = 22). Individual ROIs in the hemisphere contralateral to the dominant hand were converted to MNI space and projected onto an inflated surface. The colour code represents the number of participants with overlapping ROIs in the standard MNI space. The black contour shows the anatomical delineation of S1 used to restrict the ROI definition, based on a probabilistic atlas. **B)** Univariate activity levels (vs rest) for the three body parts (green: feet, red: hand, blue: lips) within each ROI. Only the primary body part of each ROI exhibited activity levels significantly above zero. **C)** Multivariate dissimilarities. The left plots are a multidimensional scaling (MDS) depiction of the representational dissimilarity between the three body parts (green: feet, red: hand, blue: lips) in each ROI. Ellipses indicate between-participant standard errors. The right histograms show the cross-validated dissimilarity (a.u.) observed for the three pairs of body parts in each ROI (yellow: feet-hand, cyan: feet-lips, magenta: hand-lips). Grey dots represent individual participants. Asterisks indicate a significant difference relative to zero; \**p* < 0.017; \*\*\**p* < 0.001.

### INFORMATION FROM DIFFERENT BODY PARTS IS DISTRIBUTED ACROSS S1

We first focused on the body task to assess how information from different body parts is distributed across S1. To confirm the selectivity of our independent ROIs (Fig. 1A), we extracted the average univariate activity level obtained for each body part (movement vs rest) in the contralateral hemisphere and found that each ROI was highly selective to its primary body part, showing significant activity for this body part only (one sample t-tests vs zero, primary body parts: all *t*_(21)_ ≥ 10.67, all *p* < 0.001, all *d* ≥ 2.28 95% CI [1.59 2.93]; non-primary body parts: all *t*_(21)_ ≤ 1.00, all d ≤ 0.21 95% CI [-0.14 0.57]; Fig. 1B). We then used RSA to quantify the dissimilarity between activity patterns evoked by each movement (see Supplemental Information and Fig. S3A for similar analysis using the absolute difference between univariate activity levels). One sample t-tests with Bonferroni corrected alpha levels (*α* = 0.017, corrected for the three comparisons across body parts) confirmed that the representational dissimilarities were significantly greater than zero for pairs of body parts involving the primary body part of each ROI (all *t*_(21)_ ≥ 16.21, all *p* < 0.001, all *d* ≥ 3.45 95% CI [2.50 ∞]). Interestingly, significant dissimilarities were also found for cortically remote pairs of body parts, where both body parts were non-primary to the ROI (in Leg ROI: hand-lips: *t*_(21)_ = 9.93, *p* < 0.001, *d* = 2.12 95% CI [1.46 ∞]; in Hand ROI: feet-lips: *t*_(21)_ = 6.51, *p* < 0.001, *d* = 1.39 95% CI [0.88 ∞]; in Face ROI: feet-hand: *t*_(21)_ = 10.59, *p* < 0.001, *d* = 2.26 95% CI [1.57 ∞]; Fig. 1C). Thus, despite being highly selective to their primary body part, each ROI contained robust information content about non-primary body parts. This first evidence suggests that non-primary and cortically distant body parts may contribute information to a given region of the Homunculus.

### INFORMATION FROM DIFFERENT BODY SUBPARTS IS DISTRIBUTED ACROSS S1

We next assessed how information from different body *subparts* is distributed across S1. Two tasks were used for that purpose: a face task involving bilateral movements and a finger task performed with each hand (see STAR Methods). First, to verify the selectivity of the individual ROIs used for each task, we quantified the univariate activity level obtained for each subpart (vs rest) in the contralateral ROIs. Alpha was adjusted to 0.013 and 0.01, corrected for 4 and 5 comparisons (respectively) across face and hand subparts. Activity levels (averaged across hemispheres) were significantly greater than zero for all face subparts in the Face ROI (Face ROI: all *t*_(21)_ ≥ 3.51, all *p* ≤ 0.002, all *d* ≥ 0.75 95% CI [0.34 1.14]; Fig. 2A blue) and for the five fingers in the Hand ROI (all *t*_(18)_ ≥ 7.44, all *p* < 0.001, all *d* ≥ 1.71 95% CI [1.09 2.29]; Fig. 2A red). Face subparts did not significantly activate the Hand and Leg ROIs (all *t*_(21)_ ≤ -0.86, all *d* ≤ -0.18 95% CI [-0.54 0.17]. Fig. 2A blue). For some fingers, significant positive activity levels (or trends) were found for in the Face and Leg ROIs, as shown in red in Fig. 2A (significant fingers: all *t*_(18)_ ≥ 2.70, all *p* ≤ 0.015, all *d* ≥ 0.62 95% CI [0.20 1.02]; other fingers: all *t*_(18)_ ≤ 2.47, all *p* ≥ 0.024, all *d* ≤ 0.57 95% CI [0.15 0.97]). Thus, while the face task shows high selectivity to the Face ROI, finger related activity seems to be more distributed across ROIs, though activity levels were still low on average, i.e., below the liberal 2.3 threshold usually used to threshold individual data (dotted line in Fig. 2A).

**Figure 2.**
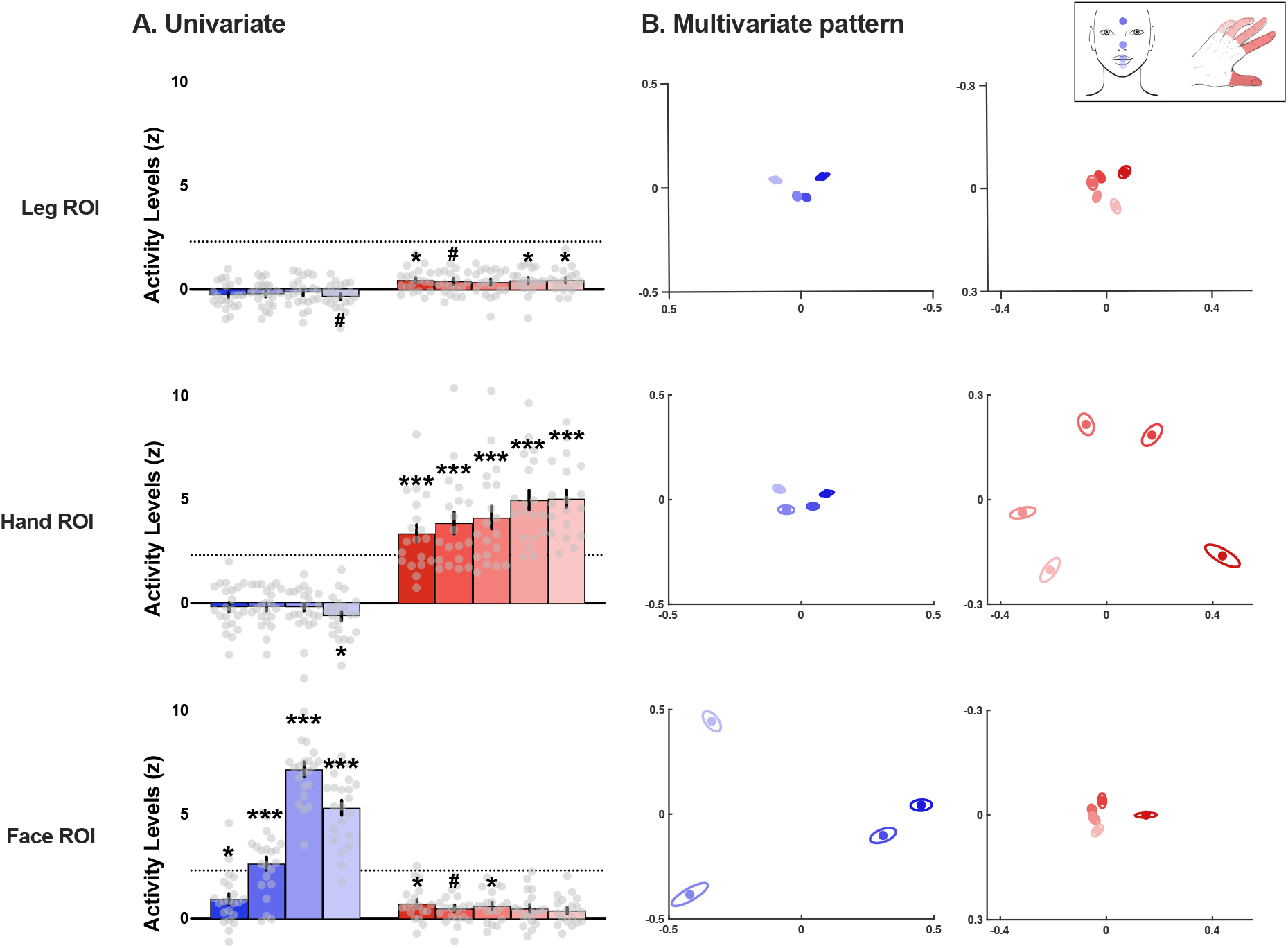
Selectivity and multivariate representational patterns of body subparts across S1 Homunculus, for face and finger tasks. **A)** Univariate activity levels (vs rest) averaged across hemispheres obtained for the different subparts of the face task (shades of blue) and of the finger task (shades of red) within each ROI. The dotted line marks the 2.3 individual threshold. **B)** MDS plots illustrating the representational structure contained in the face (shades of blue) and hand (shades of red) activity across S1 ROIs (averaged across hemispheres). The canonical hand and face representational structures are observed respectively in the Hand and Face ROIs (i.e., primary ROIs). Grey dots represent individual participants. \**p* < 0.013 and 0.010 for the face and finger tasks respectively (alpha corrected for 4 and 5 comparisons respectively); *^#^p* < 0.025 and 0.020 for the face and finger tasks respectively; \*\*\**p* < 0.001. The colour code for the respective subparts is depicted in the inset.

We then investigated the multivariate pattern of dissimilarity between the four facial subparts (Fig. 2B, blue) and the five fingers (Fig. 2B, red) across the S1 ROIs. A qualitative observation of the MDS plots (Fig. 2B) suggests that the representational structure of the face and hand, whose canonical representations are seen in their respective primary ROIs, is preserved across ROIs. This was confirmed by quantitative assessment of dissimilarities. To reduce the number of comparisons, cross-validated representational dissimilarities from different pairs of subparts (i.e., face parts or fingers) were grouped according to the subpart’s cortical neighbourhood (i.e., adjacent vs non-adjacent; see inset in Fig. 3). Alpha was adjusted to 0.025 to account for two comparisons for the adjacent and non-adjacent dissimilarities. We found that for both tasks, dissimilarities between subparts’ activity patterns were all significantly above zero regardless of their neighbourhood (Fig. 3B). This was true not only in their respective primary ROI (all *t* ≥ 12.41, all *p* < 0.001, all *d* ≥ 2.85 95% CI [1.96 ∞]) but also in remote parts of the Homunculus such as the Hand ROI or the Leg ROI for the face task (all *t*_(21)_ ≥ 5.85, all *p* < 0.001, all *d* ≥ 1.29 95% CI [0.80 ∞]; Fig. 3B blue) and the Face ROI or the Leg ROI for the finger task (all *t*_(18)_ ≥ 3.18, all *p* ≤ 0.003, all *d* ≥ 0.73 95% CI [0.29 ∞]; Fig. 3B red). These results replicate the previous observation that information about body parts is not restricted to their primary S1 area.

**Figure 3.**
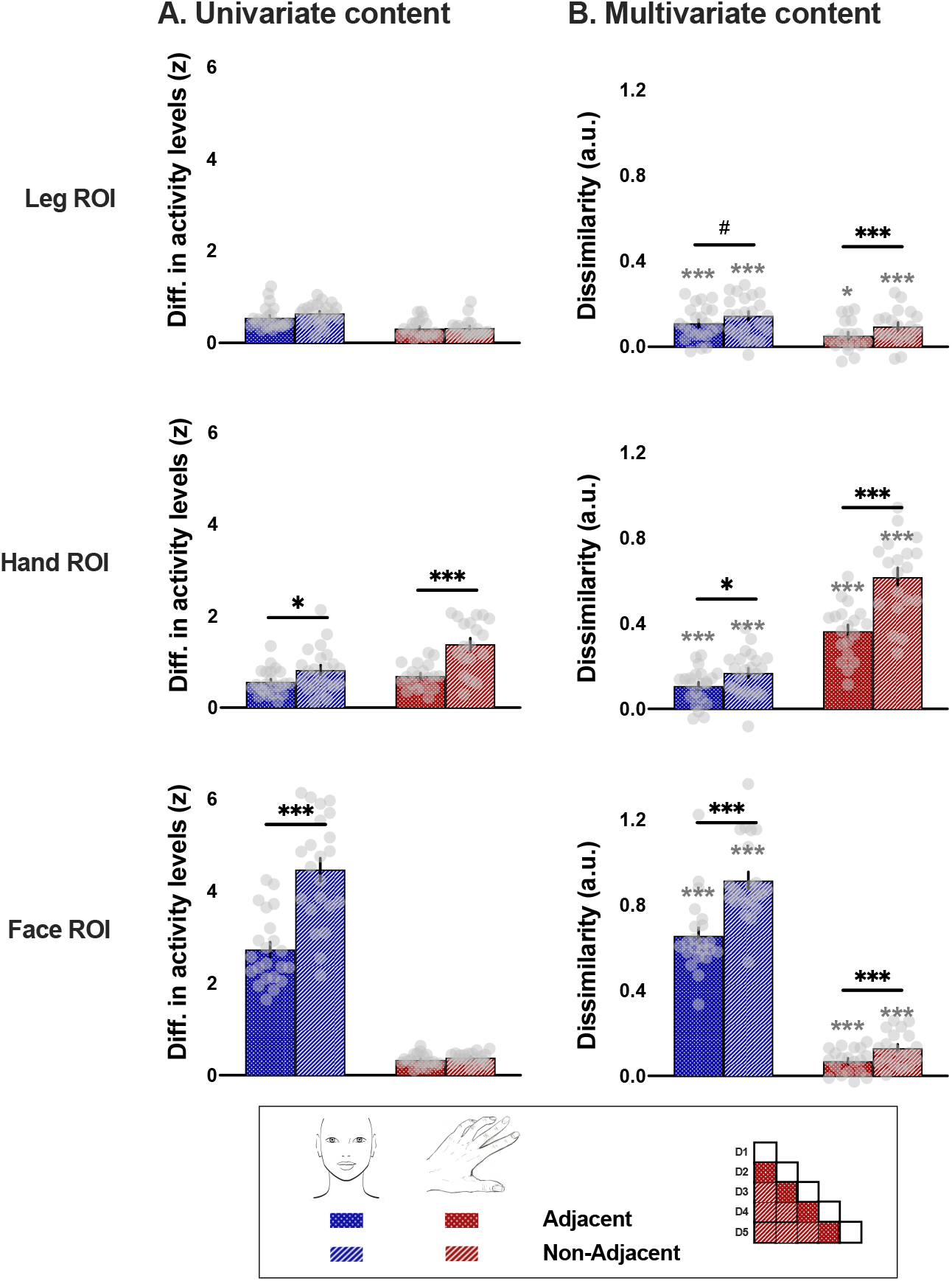
Univariate and multivariate topographic content related to body subparts across S1 Homunculus, for face and finger tasks. **A)** Univariate topographic content defined as the absolute difference between activity levels evoked by adjacent (dotted bars) and non-adjacent (hatched bars) subparts in the different ROIs for the face task (blue) and the finger task (red). **B)** Multivariate topographic content measured by the cross-validated representational dissimilarity (a.u.) between activity patterns evoked by adjacent (dotted bars) and non-adjacent (hatched bars) subparts in the different ROIs for the face task (blue) and the finger task (red). Grey dots represent individual participants. The matrix in the inset illustrates how adjacent and non-adjacent content is computed using the fingers as an example (D=digit). Black asterisks indicate a significant difference between adjacent and non-adjacent body subparts: \**p* < 0.05; *^#^p* < 0.1; \*\*\**p* < 0.001. Grey asterisks indicate a significant difference relative to zero: \**p* < 0.025; \*\*\**p* < 0.001.

### TOPOGRAPHIC FEATURES FROM DIFFERENT BODY SUBPARTS ARE DISTRIBUTED ACROSS S1

To assess whether the topographical content was preserved throughout S1, we next investigated the univariate and multivariate differences between adjacent and non-adjacent subparts across ROIs, for both the face and finger tasks. Univariate content was defined as the absolute difference between activity levels evoked by pairs of subparts in the different ROIs (see inset in Fig. 3). A significant difference between adjacent and non-adjacent univariate content was found in the primary ROI of each task (both *t* ≥ -7.51, both *p* < 0.001, both *d* ≥ -1.62 95% CI [-2.25 -0.97]; Fig. 3A). In addition, a significant topographic difference was found for the face task in the Hand ROI (*t*_(21)_ = -3.30, *p* = 0.003, *d* = -0.70 95% CI [-1.16 -0.23]), but not for other comparisons/ROIs (all *p* ≥ 0.099, all *d* ≤ -0.40 95% CI [-0.86 0.07]). Altogether, these results suggest that the univariate information content does not appear to be consistently topographically organised outside of its primary ROI.

We then compared the representational dissimilarities between adjacent and non-adjacent subparts, expecting adjacent subparts to be more similar if topographic information of body subparts is preserved across the homunculus. Similar to the univariate results a significant difference between adjacent and non-adjacent subparts was found in the primary ROIs for both tasks (both *t* ≥ -13.59, both *p* < 0.001, both *d* ≥ -2.90 95% CI [-3.85 -1.92]). Importantly, we found significant evidence for topographical content for both tasks in the non-primary ROIs (all *t* ≥ -3.41, all *p* ≤ 0.003, all *d* ≥ -0.73 95% CI [-1.19 -0.25]; Fig. 3B), with a trend found for the face task in the Leg ROI (*t*_(21)_ = -2.03, *p* = 0.055, *d* = -0.43 95% CI [-0.87 0.01]). These multivariate results reveal that topographical information content about body parts, and the hand in particular, can be observed throughout the Homunculus.

### TWO ACTIONS DONE WITH THE SAME BODY PART CAN BE DECODED IN NON-PRIMARY AREAS OF THE HOMUNCULUS

We then assessed how information from different actions done with a given body part is distributed across S1. For that purpose, we compared the squeeze and push conditions performed with each of three body parts (i.e., feet, hand and lips). Alpha was adjusted to 0.017, correcting for the three comparisons across body parts. Activity levels evoked by these actions were significantly different only when performed with the primary body part of each ROI (all *p* ≤ 0.008; Fig. 4A). No significant differences were observed in non-primary ROIs (all *p* ≥ 0.050), except for a trend for feet movements in the Hand ROI (*t*_(21)_ = -2.55, *p* = 0.019; Fig. 4A). However, multivariate representational dissimilarities were significantly greater than zero not only in primary ROIs (all *p* < 0.001, all *d* ≥ 1.00 95% CI [1.00 ∞]), but also in non-primary ROIs for hand and feet movements (all *p* ≤ 0.002, all *d* ≥ 0.69 95% CI [0.29 ∞]; except for a trend for feet movements in Face ROI: *t*_(21)_ = 2.09, *p* = 0.024, *d* = 0.45 95% CI [0.07 ∞]; Fig. 4B). These results suggest that action-related information content from the hand and the feet seems to be distributed across the Homunculus (see Methods for potential explanations for the lack of lips information).

**Figure 4.**
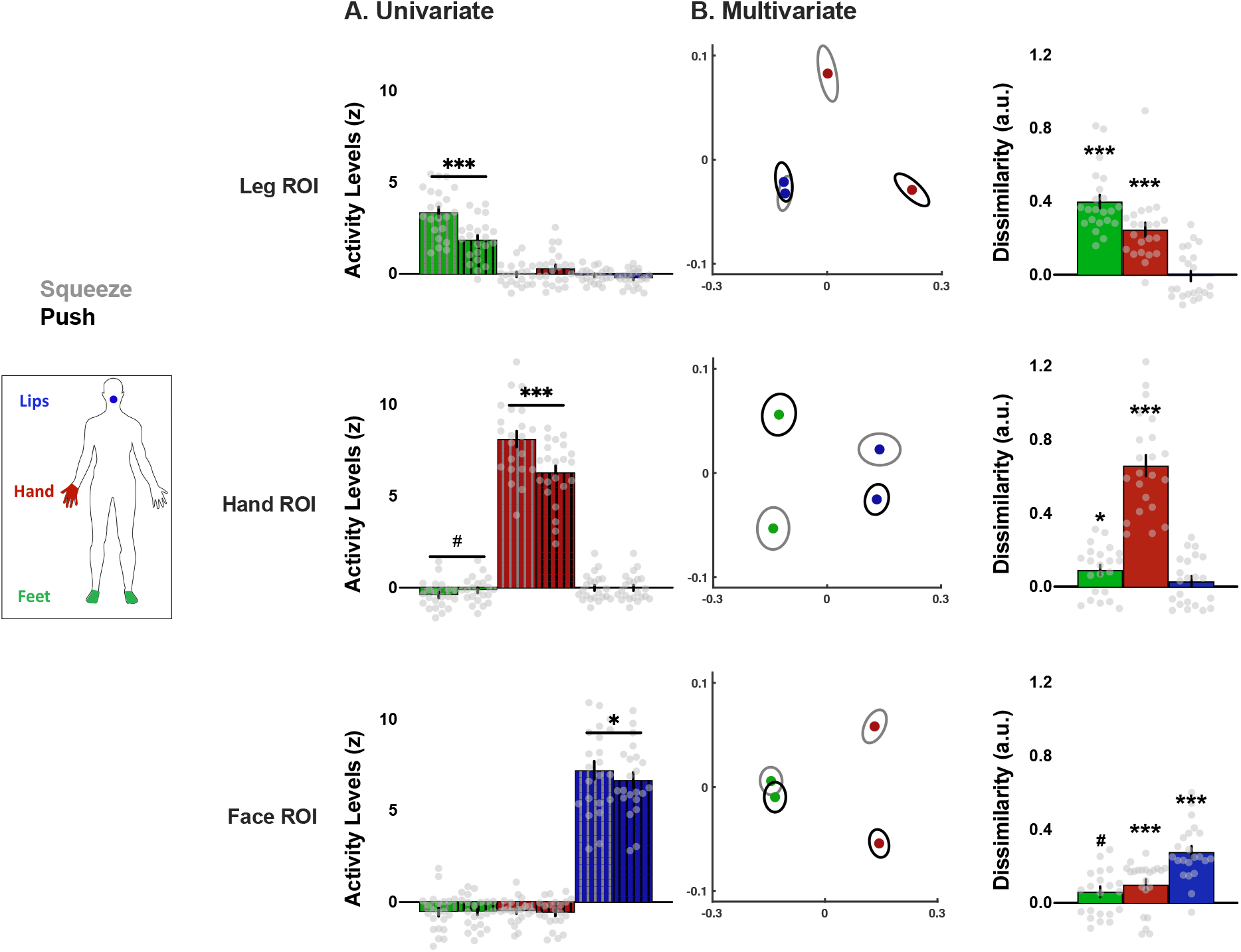
Univariate and multivariate content related to different actions performed with a given body part across S1 Homunculus. **A)** Univariate activity levels (vs rest) for the two actions (grey hatched: squeeze; black hatched: push) performed with each body part (green: feet, red: hand, blue: lips) within each ROI. **B)** Multivariate dissimilarities. The left plots are a multidimensional scaling (MDS) depiction of the representational dissimilarity between the two actions (grey ellipses: squeeze; black ellipses: push) performed with the non-primary body parts in each ROI (green: feet, red: hand, blue: lips). Ellipses indicate between-participant standard errors. The right histograms show the cross-validated dissimilarity (a.u.) observed for the two actions performed with each body part in each ROI (green: feet, red: hand, blue: lips). Grey dots represent individual participants. \**p* < 0.017; *^#^p* < 0.033; \*\*\**p* < 0.001.

### TWO ACTIONS DONE WITH THE SAME BODY PART CAN BE DECODED THROUGHOUT BA3b

Finally, we investigated the profile of action-related information content specifically in BA3b, known to show the greatest level of selectivity in S1 (Powell and Mountcastle, 1959; Martuzzi *et al*., 2014; Schellekens *et al*., 2021), alongside the univariate activity levels used to classically determine body maps (Fig. 5). To this end, BA3b’s strip was segmented into 29 bands of equal height (Fig. 5A) that were then used to calculate activity levels and multivariate dissimilarities at the individual level. Consistent with our previous ROI selectivity analysis, the highest activity levels for each body part, based on univariate analysis, lay within their independently defined primary ROI (grey shades in Fig. 5B). As we found before, multivariate dissimilarities were qualitatively apparent beyond the regions where activity levels can be observed (Fig. 5C). To reduce the number of comparisons, the peak(s) dissimilarity between two different movements with each primary body part (doted black lines in Fig. 5C) was used to test whether action dissimilarities obtained in the corresponding BA3b band using the other non-primary body parts were significantly greater than zero. Alpha was corrected to 0.025, to account for the two comparisons for the two non-primary body parts. For instance, while feet activity levels were observed solely within the first few medial bands of BA3b, dissimilarities between the two actions performed with the feet were significantly greater than zero at the two peaks observed for the hand (both *z* ≥ 208.00, both *p* ≤ 0.003, both *d* ≥ 0.64 95% CI [0.35 ∞]), but also at the peak observed for the lips (*t*_(21)_ = 3.35, *p* = 0.002, *d* = 0.71 95% CI [0.31 ∞]). Similarly, dissimilarities between the two actions performed with the hand were significantly greater than zero at the peaks observed for both the feet and lips (both *t* ≥ 2.48, both *p* ≤ 0.011, both *d* ≥ 0.53 95%CI [0.15 ∞]). These results emphasise the availability of body part information across S1. Therefore, we find that the information content about body part actions is much more widely distributed then can be inferred by delineating the univariate selectivity profiles.

**Figure 5.**
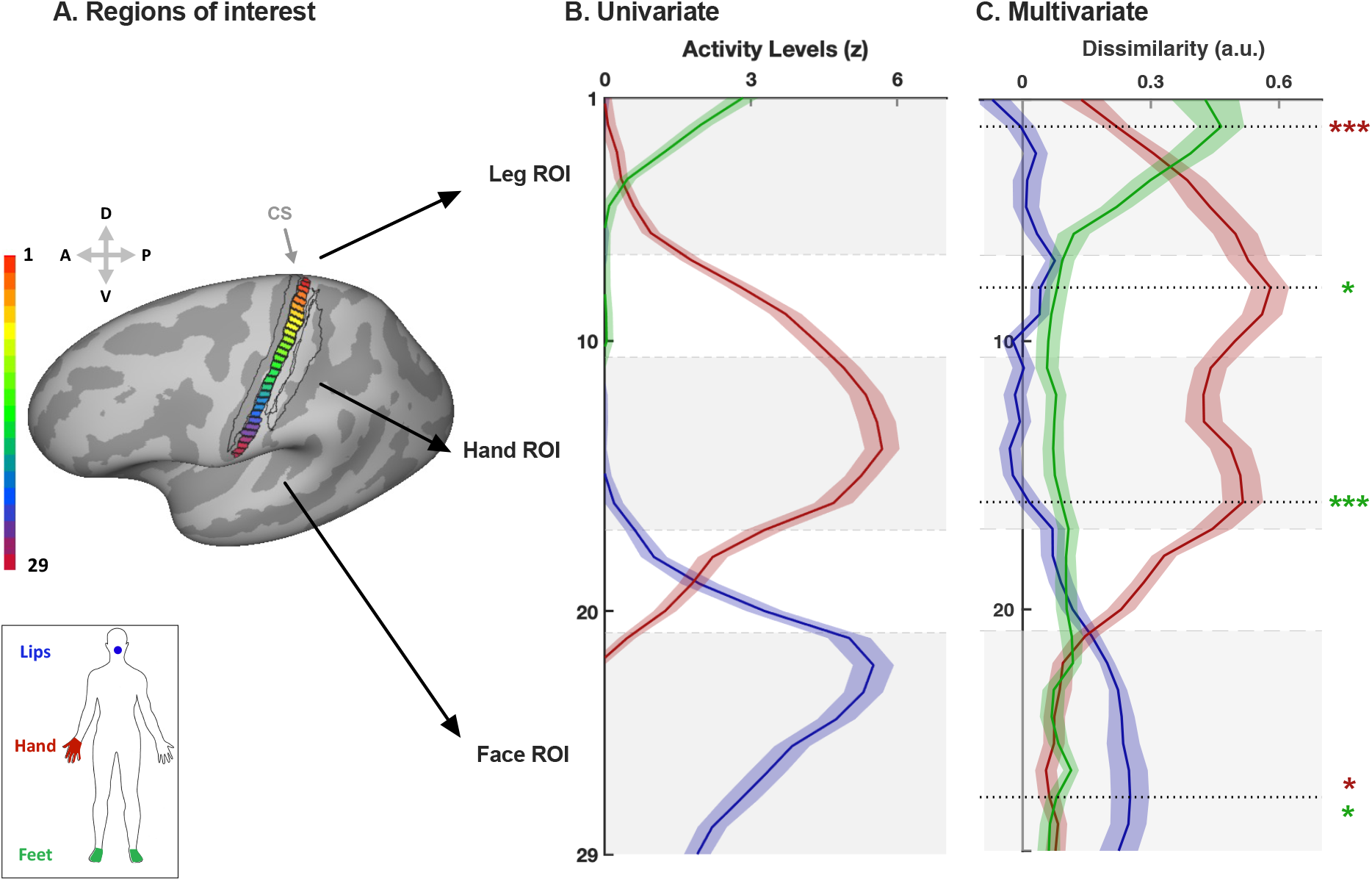
Regions of interest, selectivity, and multivariate information content related to the two actions across BA3b’s strip. **A)** Illustration of the segmentation of BA3b’s strip into 29 bands of similar height (i.e., 2.0911mm). The black outlines represent the surrounding S1 Brodmann areas, respectively BA3a, BA1 and BA2 (from left to right), based on a probabilistic atlas. The colour code represents the band number (1 to 29). CS = Central Sulcus. **B)** Univariate activity levels (vs rest) observed for the three body parts throughout BA3b strip (green: feet, red: hand, blue: lips). **C)** Multivariate cross-validated dissimilarities (a.u.) observed between the two actions (i.e., squeeze and push) for the three pairs of body parts throughout BA3b strip (green: feet, red: hand, blue: lips). The peak dissimilarity for each primary body part (doted black lines) was used to test whether dissimilarities obtained in the corresponding band of BA3b for the other non-primary body parts was significantly greater than zero; \**p* < 0.025; \*\*\**p* < 0.001. Shades around each curve represent the standard error. Grey shades in the background of each plot represent the location of our Leg, Hand and Face ROIs.

## Discussion

Due to its highly selective profile, conventional mapping procedures providing a ‘parcellated’ - all or nothing (i.e., winner-takes-all) - view over S1 have dominated our conceptualisation of its functional organisation (Roux, Djidjeli and Durand, 2018; Willoughby, Thoenes and Bolding, 2021). Consequently, alteration of map boundaries have been commonly interpreted as cortical reorganisation, with the limitations previously discussed (Muret and Makin, 2021). Using conventional univariate analyses, together with multivariate RSA, we investigated the distribution of information content underlying S1 topographic organisation. We found that S1 contained significant task-relevant information content beyond the primary area of a given body part, as defined by conventional mapping criteria. Even though, as expected from somatotopy, information content was more pronounced in primary areas, cortically distant body parts but also body subparts (e.g., fingers) could be consistently decoded throughout S1. Perhaps most strikingly, different actions performed with the hand or the feet could be decoded at remote extremities of the Homunculus. Overall, our results suggest a widespread distribution of information content across the S1 Homunculus that goes beyond what can be expected from its selectivity profile and emphasises the need to consider S1 more as a whole than as a patchwork of independent body maps. Our results also stress the need to further investigate the functional relevance of the distributed information and its potential for rehabilitation, augmentation, or brain-machine interfaces.

The widespread availability of body part information was further confirmed by focusing the analysis for decoding different movement done with the same body part along the most topographically organised sub-region of S1, BA3b. This analysis revealed that outside primary areas, information content was relatively continuous throughout BA3b. Indeed, BA3b contained relevant information about body parts and actions in a spatially continuous way throughout its strip. It is interesting to note that “boundaries” or transitions between body maps as defined by univariate data, did not seem to abruptly disrupt the distribution of information content, though the amount of information does decrease. In other words, no information loss or abrupt fluctuation were observed at functional “boundaries” between body maps, boundaries that are traditionally defined by contrasting univariate activity (e.g., between hand and face) (Kuehn *et al*., 2017). Moreover, our data suggests that functional activity and information content is not restricted or compartmentalised by anatomical septa (Fang, Jain and Kaas, 2002; Kuehn *et al*., 2017). Also, when present (i.e., for hand and feet), information content did not seem to decrease linearly with cortical distance (e.g., feet dissimilarities appeared relatively constant outside the feet area, see Fig. 5). Altogether, the widespread distribution of information content even within BA3b suggests that our spatial definition of body part representation within S1 may only reflect the “tip of the iceberg”.

In contrast to the hand and feet, lip action-related information content appeared to be more restricted to the face region (see Fig. 5). This result contrasts with the widespread topographical content we found throughout the Homunculus for the face subparts (see Fig. 3). The lack of action-related information for the lips could arise from a lower extent and richness of sensory feedbacks since contrary to the other body parts, most participants did not manipulate an object during the two lips actions (see STAR Methods). Alternatively, the lips could require less coordination with other body parts, resulting in less representational overlap. Specifically, when coordinating actions, the face is most often the recipient of targeted actions but not the supplier, contrary to the limbs. This idea is also compatible with the observation of higher resting-state functional connectivity between the hand and feet areas than between the face and the other areas in BA3b (Thomas *et al*., 2021).

In our view, our observation for widespread information content for body (sub)parts and actions across S1 are not surprising – some extent of distributed tactile information in S1 was already documented by Penfield and Boldrey, based on microstimulation (1937). We also do not believe these findings could be discarded as an fMRI artefact due to contribution of blood stealing effects (Woolsey *et al*., 1996; Harel *et al*., 2002; Devor *et al*., 2005), where local increase in blood flow also results in a decreased blood flow in the immediate surrounding areas. This is because we observe abundant information content remotely from the primary area, which likely extend beyond the spatial scale of these effects (Woolsey *et al*., 1996; Devor *et al*., 2005). In contrast, the information we detect are likely related to patterns of negative BOLD responses previously observed in S1 (Tal, Geva and Amedi, 2017), responses that were linked to neuronal activity rather than blood flow stealing effects (Shmuel *et al*., 2006; Schridde *et al*., 2008; Mullinger *et al*., 2014). Moreover, distributed information content was also detected using electrocortical recordings, where such limitations are not at play (Willett *et al*., 2020). We add to this body of previous findings by demonstrating that information is abundant in healthy participants, i.e., there is no need to recruit or consider plasticity mechanisms to observe this crossover of information across body part areas. We also demonstrate that body part information can be found across the homunculus and is not restricted to the hand area. Nevertheless, we do find that qualitatively greater sharing of information exists between the hand and the feet, as well as between the hand and the mouth. This could be driven by the topographic relationship of the primary areas (where the hand is roughly equidistant between the feet and the mouth) or due to unique functional coordination between the hand and mouth (e.g. for feeding) and the upper and lower limbs (e.g. for locomotion and balancing).

It could be argued that the use of active paradigms in the present study could result in more distributed information than would be obtained with a passive tactile paradigm. While this needs to be investigated, it is important to note that recent studies using similar multivariate analyses in the hand area showed that the representational multivariate structure, as well as the univariate topographic map, were comparable between active paradigm and passively applied tactile stimulation to individual fingers (Berlot *et al*., 2019; Sanders *et al*., 2019). Comparable decoding of hand gestures or postures was also previously reported in humans using electrocorticography, with similar decoding in S1 (Chestek *et al*., 2013; Branco *et al*., 2017; Li *et al*., 2017), and in M1 (Branco *et al*., 2017). In addition, recent studies showed that finger movements and effector information can be decoded in S1 during motor planning, well before movement execution (Ariani, Pruszynski and Diedrichsen, 2020; Gale, Flanagan and Gallivan, 2021). Thus, an active paradigm allows us to take full advantage of sensorimotor information, relevant for motor planning (Sun *et al*., 2015) and encompass signal arising from the efference copy (London and Miller, 2013). This abundance of tactile, proprioceptive, and even cognitive (Meftah, Shenasa and Chapman, 2002) inputs is crucial for S1 function and as such our active paradigm is arguably more appropriate to investigate ecological representational motifs.

Another important consideration is that we found this distributed information because decoding techniques are too sensitive. For example, the existence of distributed tactile information outside of the sensorimotor cortex was recently detected in neurons as far as in the rodent primary visual cortex (Enander *et al*., 2019). It is important to acknowledge that decoding does not necessarily mean the brain is actually using this information (i.e., functionally represented). Indeed, microsimulation in the S1 hand area most often elicits sensations on the patients’ hand (Roux, Djidjeli and Durand, 2018) and stroke in the M1 hand area results in motor impairment of the hand (Darling *et al*., 2016). It is important to clarify that we do not negate the notion of a primary function for a given S1 region. But it is also worth noting that these clinical observations in humans are relatively crude (Richards, Malouin and Nadeau, 2015), and do not contradict the notion that the latent activity, which comprises the information content we decoded, might also be functionally relevant. For example, it has been suggested that latent activity in M1 could contribute to inhibit movement in other body parts not involved in the task (Zeharia *et al*., 2012), or to afford better motor coordination across body parts (Graziano, Taylor and Moore, 2002). Similarly, latent activity in S1 could serve a role for predicting and encoding whole-body sensory feedback expected and perceived during actions involving multiple body parts. Distributing the content of information throughout S1 could allow for an increased number of combinations and patterns throughout body parts (Hoffmann *et al*., 2018), which might be more ecologically-relevant, considering that we rarely use body parts independently from each other. In other words, this distributed information could provide a way to support coordinated movements between body parts and to give context to their resulting sensory inputs in a coherent manner.

Even if the distributed information underlying the traditional Homunculus may not serve a functional role under normal circumstances, it could represent an underlying “scaffolding” for plasticity to take place, such as following congenital (Hahamy *et al*., 2017) or acquired (Pons *et al*., 1991) deprivation. For example, latent activity could be harnessed to restore the deprived primary function by potentiating any residual, now latent, activity (Qi, Kaas and Reed, 2014). This idea could open new perspectives for rehabilitative strategies. Moreover, even if not functionally valuable, this information could be exploited for brain-machine interfaces (Willett *et al*., 2020), where specific parts of the homunculus might not be as directly accessible (e.g., the medial foot area). Finally, the distribution of information across the homunculus, and redundancy of information that it might entail, could prove particularly useful for solving the issue of ‘resource reallocation’ that augmentation techniques are currently facing (Dominijanni *et al*., 2021).

To conclude, our results suggest that information in S1 might be a lot more distributed than selectivity profiles and winner-takes-all mapping approaches lead us to presume. While the functional consequences of this widespread information need to be further investigated, it reveals yet unexplored underlying information contents that could be harnessed for rehabilitation, augmentation or brain-machine interfaces.

## Acknowledgements

This work was supported by an ERC Starting Grant (715022 EmbodiedTech) and a Wellcome Trust Senior Research Fellowship (215575/Z/19/Z), awarded to TRM. We thank Arabella Bouzigues and Maria Kromm for their substantial help in terms of recruitment and data collection, we also thank Adriana Zainurin, Esther Teo, Christine Tan and Mathew Kollamkulam for help with data collection.

## Author contributions

D.M. and T.R.M. conceived the study. D.C. designed the objects. D.M. collected the body and face datasets, P.K. and D.C. collected the finger dataset. V.R. pre-processed the face dataset, D.M. pre-processed the body dataset and analysed the body and face datasets, P.K. pre-processed and analysed the finger dataset. D.M. and T.R.M. wrote the manuscript with inputs from all co-authors. T.R.M. secured funding.

## STAR Methods

### RESOURCE AVAILABILITY

#### Lead contact

For further information and requests for resources should be directed to and will be fulfilled by the lead contacts, Dollyane Muret and Tamar R. Makin.

#### Materials availability

The study did not generate new materials.

#### Data and code availability

All data will be made available prior to publication in a public depository (https://osf.io).

### EXPERIMENTAL MODEL AND SUBJECT DETAILS

Twenty-two healthy volunteers [mean age = 45.55 ± 9.47 (SD) years; 10 women; 6 left-handed] took part in the body and face tasks and a further nineteen healthy volunteers [mean age = 23.16 ± 4.34 (SD) years; 11 women; all right-handed] took part in the finger task. To account for age-related differences, age was added as a covariate in statistical analyses. Participants reported no sensorimotor disorders and had no counterindications for magnetic resonance imaging. All participants gave written informed consent before participating. The protocols were approved by the NHS National Research Ethics Service approval (18/LO/0474) for the body and face tasks and UCL Research Ethics Committee (REC: 12921/001) for the finger task and were performed in accordance with the Declaration of Helsinki. The face and hand datasets were recently used for other purposes (Kieliba *et al*., 2021; Root *et al*., 2021).

## METHODS DETAILS

### Scanning procedures

Each dataset comprised three or four functional task-related block-design runs, a functional localiser, a structural scan and field maps.

#### Body and face tasks

The two tasks were performed within the same experimental session. Prior to entering the scanner room, participants were thoroughly instructed, and all movements were practiced in front of the experimenter to ensure they were performed correctly. For the body task, participants were instructed to perform one of two actions (i.e., squeeze or push; Fig. S1) with one of three different body parts (i.e., feet, dominant hand and lips), resulting in a total of six conditions. Two additional conditions involving the non-dominant arm were also included but will not be described in the main text since we focus on the hemisphere contralateral to each body part. See Supplemental Information for similar analyses and results in the non-dominant hemisphere for the body task (Fig. S2 and S5). For the face task, the full details of the procedures and acquisition parameters can be found in Root et al, 2021. In short, participants were instructed to perform one of four movements: raise the eyebrows (i.e., forehead), flare nostrils (i.e., nose), puckering lips (i.e., lips), and tap the tongue to the roof of the mouth (i.e., tongue). Two additional conditions involving the left and right thumbs were also included but will not be further described as they were not included in the main analysis.

For both tasks, instructions and pace were provided visually via a screen, resulting in 5 cycles of movement per 8 sec block. In addition, each movement block was repeated 4 times per run, which also comprised 5 blocks of rest used as baseline. Conditions were pseudo-randomly distributed, such that each condition was equally preceded by all other conditions. Three and four functional runs were acquired for the face and body tasks, respectively. To confirm that appropriate movements were made at the instructed times, task performance was visually monitored online.

#### Finger task

The full details of the procedures and acquisition parameters can be found in Kieliba et al, 2021. In short, participants performed an active finger tapping task using a button box. Each finger movement was repeated at 1Hz over a period of 9s per block, with 4 blocks per finger per run and 4 runs in total. Instructions and pace were provided visually, and task performance was monitored online.

#### Functional localiser

Participants were visually instructed to move one of five body parts: right or left hand (open/closing the fingers), right or left toes (wiggling the toes) or lips (puckering the lips). Movements were repeated at a constant instructed pace for a period of 12s, interleaved with 12s of rest. Each condition was repeated 4 times in a pseudo-random order. Here again, participants practiced the movements before entering the scanner and task performance was visually monitored online.

### MRI data acquisition

MRI images were acquired using a 3T Prisma MRI scanner (Siemens, Erlangen, Germany) with a 32-channel head coil. Functional data were obtained using a multiband T2*-weighted pulse sequence with a between-slice acceleration factor of 4 and no in-slice acceleration. The following acquisition parameters were used: TR = 1450 ms; TE = 35 ms; flip angle =70°; voxel size = 2 mm isotropic; imaging matrix = 106 x 106; FOV = 212 mm. 72 slices were oriented in the transversal plane covering the entire brain. A total of 216, 172 and 346 volumes were collected per participant for each run of the body, face and finger tasks respectively. Field-maps were acquired for field unwarping. A T1-weighted sequence (MPRAGE, TR = 2530 ms; TE = 3.34 ms; flip angle = 7°; voxel size = 1 mm isotropic) was used to obtain anatomical images.

## QUANTIFICATION AND STATISTICAL ANALYSIS

MRI analysis was implemented using tools from FSL (Smith *et al*., 2004; Jenkinson *et al*., 2012), Connectome Workbench software (humanconnectome.org) in combination with bash and Matlab scripts (version R2016a), both developed in-house and as part of the RSA Toolbox (Nili *et al*., 2014). Cortical surface reconstructions were produced using FreeSurfer (version 7.1.1; Dale, Fischl, and Sereno 1999; Fischl, Liu, and Dale 2001, freesurfer.net).

### fMRI pre-processing

Functional data was pre-processed in FSL-FEAT (version 6.00) and included the following steps: motion correction using MCFLIRT (Jenkinson *et al*., 2002); brain extraction using BET (Smith, 2002); high-pass temporal filtering with a cut-off of 150s, 119s and 150s for the body, face and finger tasks respectively and 280s for the functional localiser; and finally spatial smoothing using a Gaussian kernel with a full width at half maximum of 3mm for the three tasks, and 5mm for the functional localiser. Field maps were used for distortion correction of the functional data from the body and face tasks and the functional localiser collected for these participants. For each participant, a midspace between the different functional runs of each task was calculated, i.e., the average space in which the images are minimally reorientated. Each functional run was then aligned to the midspace and registered to each individual structural T1 scan using FMRIB’s Linear Image Registration Tool (FLIRT), optimised using Boundary-Based Registration (Greve and Fischl, 2009). Where specified, functional and structural data were transformed to MNI152 space using FMRIB’s Nonlinear Registration Tool (FNIRT).

### fMRI low-level analysis

Voxel-wise General Linear Model (GLM) was applied to the data using FEAT to obtain statistical parametric maps for each movement. For each task, the design comprised a regressor of interest for each movement convolved with a double-gamma hemodynamic response function (Friston *et al*., 1998) and its temporal derivative. The six motion parameters were included as regressors of no interest. Large head movements between volumes (> 0.9 mm for body and face tasks, > 1 mm for finger task) were defined as motion outliers and regressed out, by adding additional regressors of no interest to the GLM [body task: mean proportion of volumes excluded = 0.45 ± 0.76 (SD) %; face task: mean proportion of volumes excluded = 0.36 ± 0.67 (SD) %; finger task: mean proportion of volumes excluded = 0.32 ± 0.77 (SD) %].

For each task, a contrast relative to rest was set up for each movement, resulting in 8 contrasts for the body task (i.e., lip, dominant hand, non-dominant arm, right/left foot x Squeeze or Push, each vs rest), in 6 contrasts for the face task (i.e., forehead, nose, lips, tongue; and left/right thumb, not used here), and in 12 contrasts for the finger task (i.e., each digit of each hand; and feet and lips, not used here). The estimates from the total number of functional runs for each task (3 for face task, 4 for body and finger tasks) were then averaged voxel-wise at the individual level using fixed effects model. For the face task, each estimates’ average was masked prior to cluster formation with a sensorimotor mask, defined as the precentral and postcentral gyrus from the Harvard Cortical Atlas. The sensorimotor mask was registered to the individuals structural scan using an inversion of the nonlinear registration by FNIRT.

For the functional localiser, each condition (i.e., right/left hand, right/left toes, lips) was contrasted against all other conditions to identify the most selective voxels. The activity patterns associated with these five contrasts were then registered to the structural space of each individual and to the functional space of each task using FLIRT to define regions of interest.

### Definition of regions of Interest (ROIs)

Since we were interested in investigating the information content of highly selective regions across the S1 homunculus, we used the functional localiser to select highly selective voxels to toe, hand and lip movements within anatomical S1 ROIs. The functional ROI was restricted by anatomical criteria, as detailed below. Although M1 is expected to be largely activated during each movement, M1 topography tends to be less well-defined and thus information content more widespread (Schieber, 2001; Graziano and Aflalo, 2007). We therefore primarily focus on the more topographically selective S1, though we wish to note that marginal contribution from M1 may have affected our S1 activity profiles due to their spatial proximity, the probabilistic nature of our anatomical ROIs and spatial smoothing of the data.

First, S1 was defined on a template surface using probabilistic cytoarchitectonic maps, by selecting nodes for Brodmann areas (BAs) 3a, 3b, 1 and 2 (Wiestler and Diedrichsen, 2013). This S1 anatomical mask was then split into three anatomical sub-regions. A node approximately 1cm below and above the hand knob was defined as an anatomical hand sub-region. Note that this criterion defined a more conservative hand region than was done in previous work (Wiestler and Diedrichsen, 2013; Wesselink *et al*., 2019; Kieliba *et al*., 2021). A gap of 1cm was then defined above and below this anatomical hand sub-region, and the remaining medial and lateral parts of S1 were used as the other two anatomical sub-regions.

Structural T1-weighted images were used to reconstruct the pial and white-grey matter surfaces using Freesurfer. Surface co-registration across hemispheres was done using spherical alignment. The three anatomical S1 sub-regions were then projected into the individual brains via the reconstructed individual anatomical surfaces. To exclude any possible contribution from neighbouring more integrative regions that contain information from multiple body parts, we further trimmed in participant’s structural space: i) the medial sub-region by removing any overlap with BA5L and BA5M, and ii) the lateral sub-region by removing any overlap with S2. BA5L, BA5M and S2 were defined in MNI152 space using the Juelich Histological Atlas thresholded at 25% maximum probability (Wiech *et al*., 2014). BA5L, BA5M and S2 were then registered to participants’ structural space using an inversion of the nonlinear registration carried about by FNIRT and used to trim our anatomical sub-regions.

These trimmed anatomical sub-regions were then registered to functional space of each task using FLIRT and used to mask the functional localiser contrasts. The medial S1 sub-region was used to mask the toe contrasts, the central hand sub-region to mask the hand contrasts and the lateral S1 sub-region to mask the lip contrast. Within each of these anatomical sub-regions, we then selected the 50 most activated voxels for the corresponding contrasts (all contrasts vs all other body parts, see section ‘fMRI low-level analysis’). This provided us with the most selective Leg, Hand and Face ROIs for each individual, while ensuring the same ROI size across participants and regions.

### Univariate analysis

The z statistic time series from the 50 voxels of each ROI obtained for each movement were extracted and averaged. These averaged values were used to assess the selectivity of our ROIs. Univariate information content was defined as the absolute difference between the averaged univariate activity evoked by two movements in a given ROI. For the body task, the two absolute differences obtained between pairs of body parts when performing the same action (i.e., squeeze or push) were averaged to define an overall difference between body parts. For the face and finger tasks, since face movements evoked bilateral activity and finger movements were performed with each hand, and since no major differences were observed across hemispheres (see Supplemental Information), absolute differences from the two hemispheres were averaged within participants. To further reduce the number of comparisons while still assessing the topographical content, absolute difference from different pairs of subparts (i.e., face parts or fingers) were grouped according to the subpart’s cortical neighbourhood (i.e., adjacent vs non-adjacent).

### Multivariate representational similarity analysis

Representational Similarity Analysis (RSA; Nili et al. 2014) was used to assess the multivariate relationship between the contralateral activity patterns generated by each movement. The dissimilarity between activity patterns within each S1 ROI (i.e., Leg, Hand and Face) was computed at the individual level for each pair of movements using cross-validated squared Mahalanobis distance (Walther *et al*., 2016). Multidimensional noise normalisation was used to increase reliability of distance estimates (noisier voxels are down-weighted), based on the voxel’s covariance matrix calculated from the GLM residuals. Due to cross-validation, the expected value of the distance is zero (or negative) if two patterns are not statistically different from each other, and significantly greater than zero if the two representational patterns are different (Diedrichsen, Provost and Zareamoghaddam, 2016). Larger distances for movement pairs therefore suggest greater information content. The resulting representational pairwise distances (i.e., 8 for the body task, 6 for the face task and 10 for the finger task) were extracted. For the body task, the dissimilarities obtained between pairs of body parts when performing similar actions (e.g., dissimilarity between lip squeeze and feet squeeze) were averaged across actions (e.g., previous example averaged with dissimilarity between lip push and feet push) to define overall dissimilarity between body parts. For the face and finger tasks, since face movements evoked bilateral activity and finger movements were performed with each hand, and since no major significant differences were observed between hemispheres (see Supplemental Information), dissimilarities from the two hemispheres were averaged within individual participants. To further reduce the number of comparisons while still assessing the topographical content, dissimilarities from different pairs of subparts (i.e., face parts or fingers) were grouped according to the subpart’s cortical neighbourhood (i.e., adjacent vs non-adjacent). Multidimensional scaling (MDS) was used to project the higher-dimensional representational dissimilarity matrices into lower-dimensional space, whilst preserving pairwise dissimilarities, for visualisation purposes only. Analysis was conducted on an adapted version of the RSA Toolbox in MATLAB (Nili *et al*., 2014), customised for FSL (Wesselink and Maimon-Mor, 2018).

### BA3b analysis

The same analyses as the ones performed in the S1 ROIs described above were performed throughout BA3b’s strip. BA3b was defined on the same template surface as S1 using probabilistic cytoarchitectonic maps, by selecting the nodes showing at least 50% maximum probability for BA3b (Wiestler and Diedrichsen, 2013). BA3b’s strip was then segmented into 30 bands, each 2.09mm high in the medio-lateral direction (see Fig. 5A). Similar to the S1 ROIs, these bands were then projected into the individual brains via the reconstructed individual anatomical surfaces and registered to participants’ functional space of each task using FLIRT. Univariate and multivariate analyses were then repeated in each of these bands. Note that the most medial band contained very few and discontinuous voxels that prevented from getting reliable RSA dissimilarities, and was thus excluded from further analysis.

### Statistical analysis

Two-tailed one-sample t-tests versus zero were used to assess significant activity levels in each ROIs. Alpha levels were Bonferroni corrected for the number of tests performed across conditions within each ROI (i.e., alpha = 0.017 corrected for three comparisons for the body task, alpha = 0.013 corrected for four comparisons for the face task and alpha = 0.010 corrected for five comparisons for the finger task). Since negative dissimilarity measures represent noise levels, one-tailed one-sample t-tests versus zero were used to test the significance of representational dissimilarities as well as absolute differences in activity levels in each ROIs. Here again, alpha levels were Bonferroni corrected for the number of tests performed within each ROI (i.e., corrected for three comparisons for the body and action dissimilarities, and for two comparisons for the adjacent and non-adjacent dissimilarities for the face and finger tasks). Paired t-tests were used to compare adjacent and non-adjacent conditions for the face and finger tasks. In each case, a trend was defined when *p* values were inferior to twice the corrected alpha level. Data normality was assessed using Shapiro-Wilk test. Effect sizes were computed using Cohen’s *d* (Cohen, 1988), and based on benchmarks suggested by Cohen, a large effect size was defined as greater than 0.8. Wilcoxon signed-rank t-tests were used when data violated normality assumptions. Two three-way rmANOVAs with the factors Hemisphere, ROI and Subpart were applied to univariate activity levels from the face and finger tasks to assess Hemisphere effect or interaction. Since no significant main effect or interaction were observed (see Supplemental Information), univariate and multivariate data from each hemisphere were averaged. Greenhouse-Geisser correction was applied when data did not follow sphericity assumption.

### Group-level ROI visualisations

S1 ROIs of each participant were projected to MNI152 space using the nonlinear registration carried about by FNIRT. Participant information regarding hand dominance were used to sagittal-flip data, such that the ROIs contralateral to the dominant hand were always represented in the left hemisphere. ROIs of all participants were then concatenated into a single volume to produce a consistency map (i.e., how many participants have their ROIs overlapping in the MNI space). Resulting consistency maps were then projected to a group cortical surface (Glasser et al., 2016) using Connectome Workbench (v1.4.2) (see Fig. 1 and S2 for body task; see Fig. S4 for face and finger tasks).

## Supplemental Information

**Figure S1.**
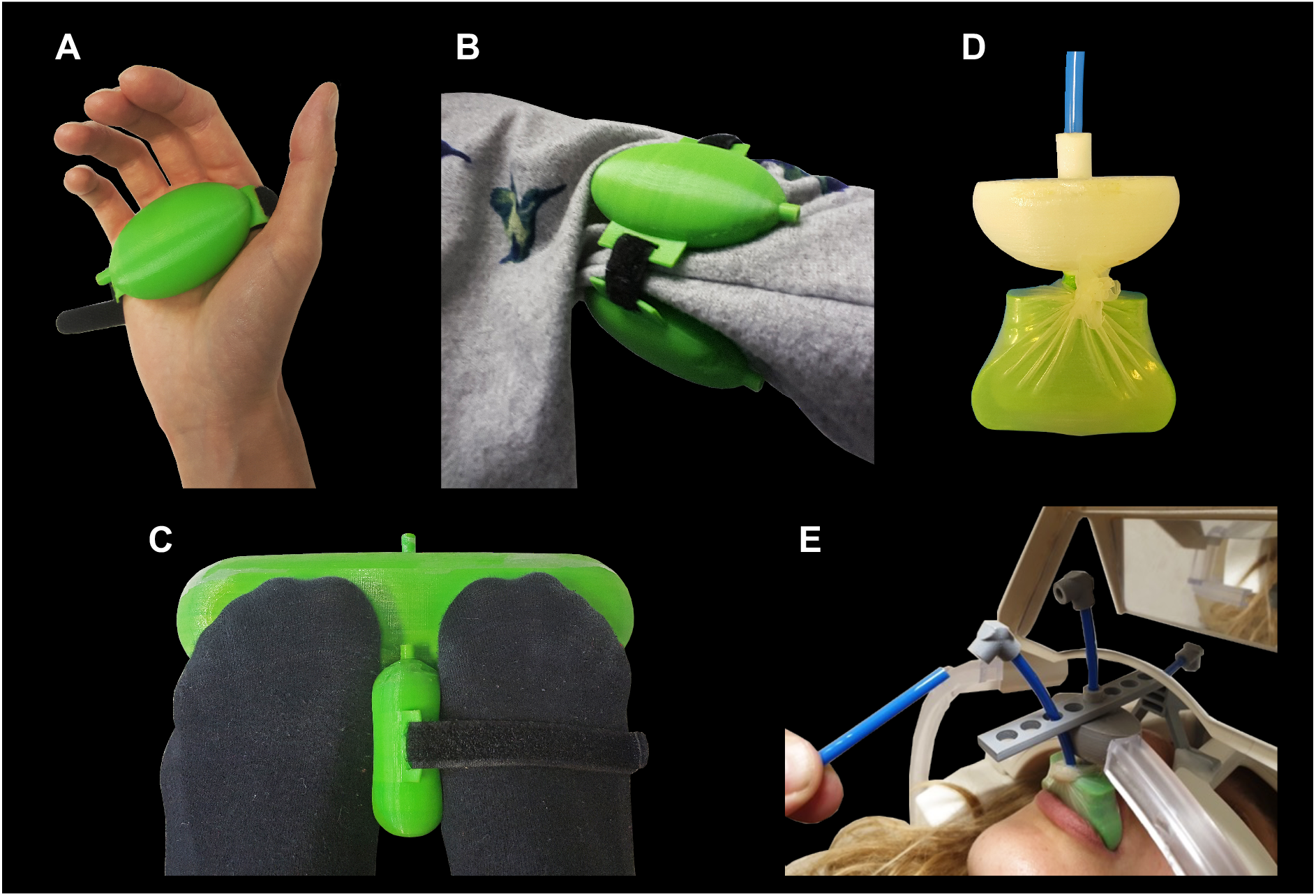
Objects used in the body task: Silicon 3D printed objects (NinjaFlex© 3D printing filament) were placed and secured with straps in the participants dominant hand (A), around the non-dominant arm (B), between the feet (secured at the level of their right metatarsophalangeal joints), and behind their toes (strapped to a footrest secured on the bed behind their feet) (C). Participants were instructed to either squeeze (i.e., closing the fingers for the hand, against the torso for the arm and between the feet) or push (i.e., against the bed for hand and arm, and against the footrest for the feet) the object with each body part. Three participants also had an object placed in between their lips (D), and secured on a fully adjustable frame, fixed on the coil (E), object that could be either squeezed or pushed. The other participants were asked to either purse their lips (equivalent to squeezing) or blow kisses (equivalent to pushing). A plastic tube was connecting each of the objects to a transducer system in the control room, allowing us to monitor and quantify the amount of pressure exerted.

### INFORMATION FROM DIFFERENT BODY PARTS IS DISTRIBUTED ACROSS S1 – non-dominant hemisphere

In agreement with what was found in the dominant hemisphere (i.e., contralateral to the dominant hand), ROIs in the non-dominant hemisphere (Fig. S2A) were highly selective to their primary body parts, showing significant activity for this body part only (Fig. S2B, primary body parts: all *t* ≥ 6.50, all *p* < 0.001, all *d* ≥ 1.39 CI [0.88 1.87]; non-primary body parts: all *t* ≤ 1.20, all *d* ≤ 0.25 CI [-0.10 0.61]). Using RSA to quantify the dissimilarity between activity patterns evoked by each movement, we found here again dissimilarities significantly greater than zero not only for pairs of body parts involving the primary body part of each ROI (all *t* ≥ 13.01, all *p* < 0.001, all *d* ≥ 2.77 95% CI [1.97 ∞]), but also for pairs of body parts non-primary to the ROI (all *t* ≥ 5.84, all *p* < 0.001, all *d* ≥ 1.25 95% CI [0.76 ∞]; Fig. S2C).

**Figure S2.**
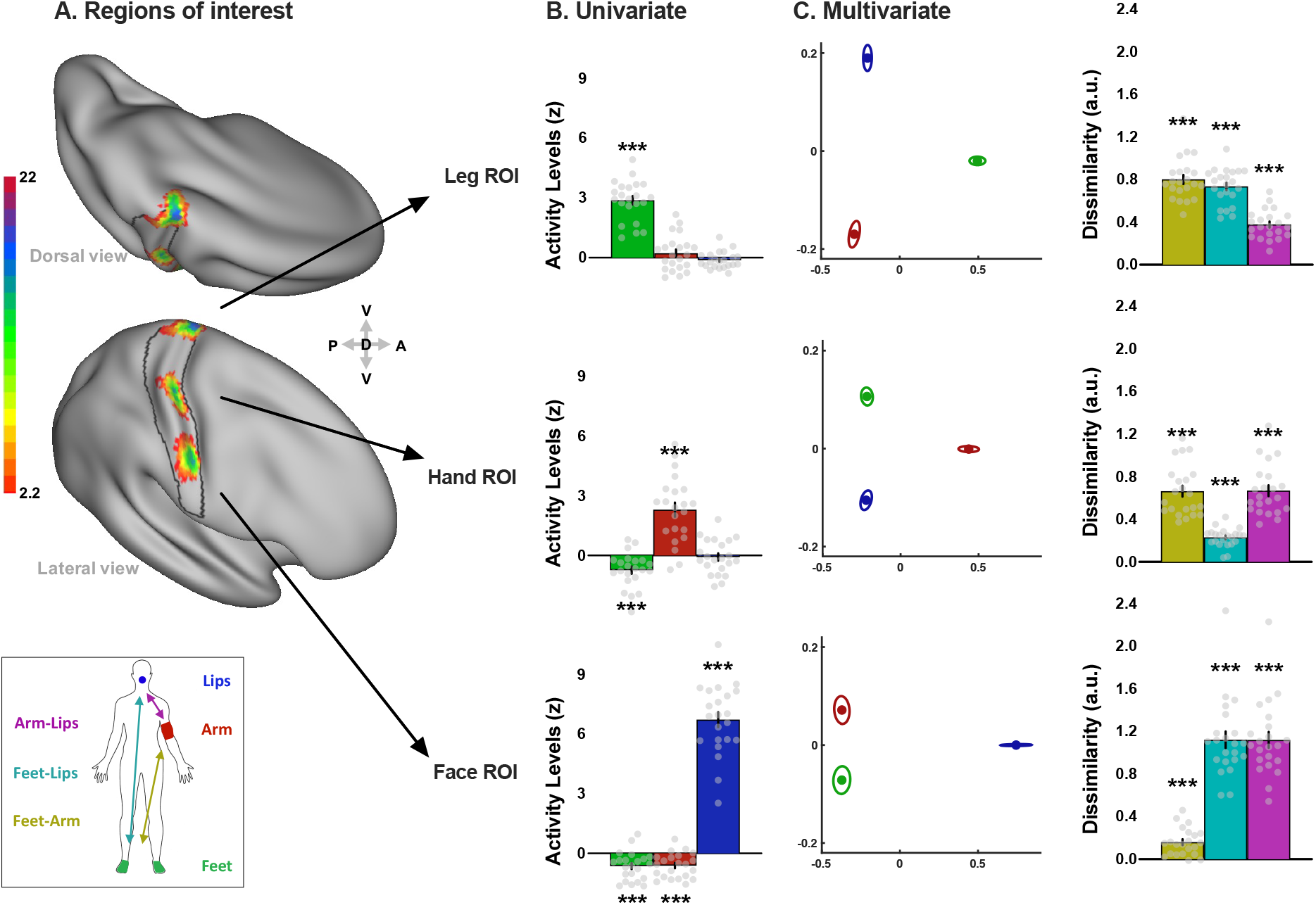
Regions of interest, selectivity, and multivariate information content about body parts across S1 Homunculus in the non-dominant hemisphere. **A)** Consistency maps across participants of the S1 regions of interest (ROIs) for the body task (n = 22). All figure annotations are as denoted in Figure 1. Asterisks indicate a significant difference relative to zero; \*\*\**p* < 0.001. We next quantified the absolute difference between the univariate activity levels observed for each pair of body parts. In each hemisphere absolute differences were significantly greater than zero not only for pairs of body parts involving the primary body part of each ROI (all *t*_(21)_ ≥ 7.96, all *p* < 0.001, all *d* ≥ 1.70 95% CI [1.13 ∞]), but also for pairs of body parts that were non-primary to the ROI (all *t*_(21)_ ≥ 5.18, all *p* < 0.001, all *d* ≥ 1.10 95% CI [0.65 ∞]; Fig. S3).

**Figure S3.**
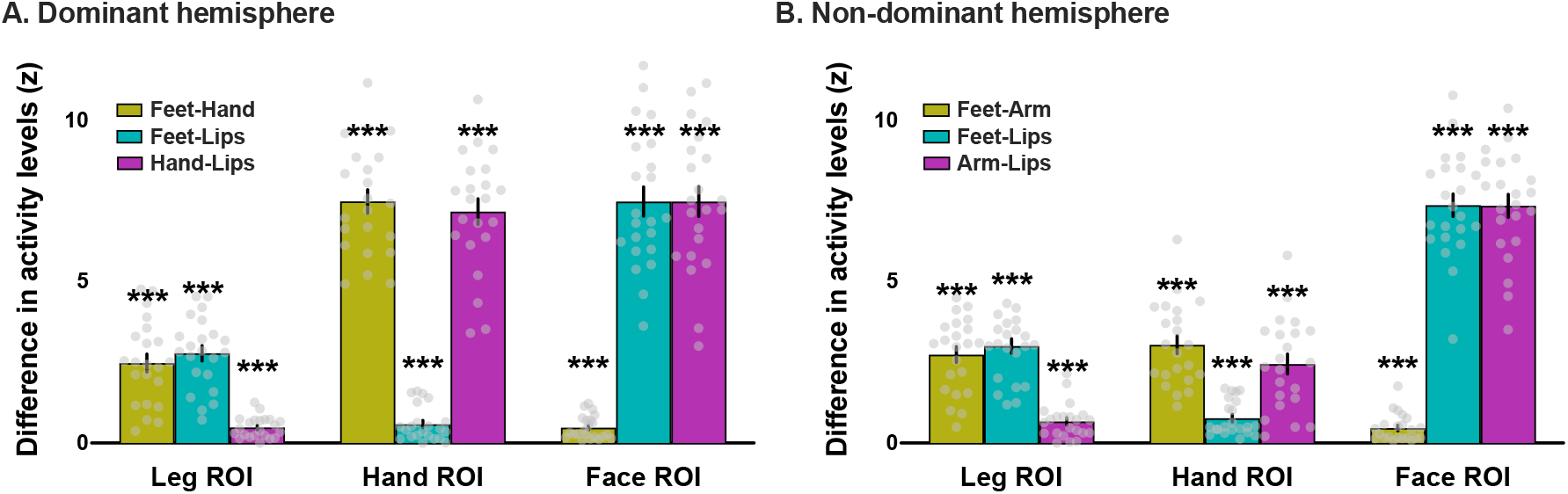
Univariate information content about body parts across the S1 Homunculus. Absolute differences in activity levels between activity levels observed for each pair of body parts (yellow: feet-hand/arm, cyan: feet-lips, magenta: hand/arm-lips) across S1 ROIs in the hemisphere contralateral to the **A)** dominant hand or **B)** non-dominant arm. Grey dots represent individual participants. \*\*\**p* < 0.001.

### INFORMATION FROM DIFFERENT BODY SUBPARTS IS DISTRIBUTED ACROSS S1

To limit the number of tests performed and since these tasks involved both body sides in a symmetrical way, we assessed whether we could pull data from the two hemispheres together by testing the presence of a main effect or interaction with the factor Hemisphere at the univariate level. A three-way repeated measures ANOVA with the factors Hemisphere, ROI and Face subparts revealed no main effect of Hemisphere (*F*_(1,21)_ = 1.30, *p* = 0.267) and no interactions with Hemisphere (Hemi*ROIs: *F*_(2,42)_ = 0.17, *p* = 0.846; Hemi*Face subparts: *F*_(2.06,43.32)_ = 0.95, *p* = 0.397) except for a triple interaction Hemi*ROIs*Face subparts: *F*_(3.12,65.49)_ = 3.06, *p* = 0.033) revealing a significant difference between hemispheres for the lips (*z*_(21)_ = 45.00, *p* = 0.007) and for the nose in the Leg ROI only (*t*_(21)_ = -2.78, *p* = 0.011). A similar analysis performed on the finger task revealed also no main effect of Hemisphere (*F*_(1,18)_ = 1.17, *p* = 0.293) and no interactions with Hemisphere (Hemi*ROI: *F*_(1.37,24.68)_ = 1.20, *p* = 0.30; Hemi*Finger: *F*_(2.88,51.77)_ = 0.86, *p* = 0.465; Hemi*ROI*Finger: *F*_(3.15,56.65)_ = 0.91, *p* = 0.444). We thus decided to average data from the two hemispheres for both tasks. See Figure S4 for the consistency across individuals of the ROIs used for each task.

**Figure S4.**
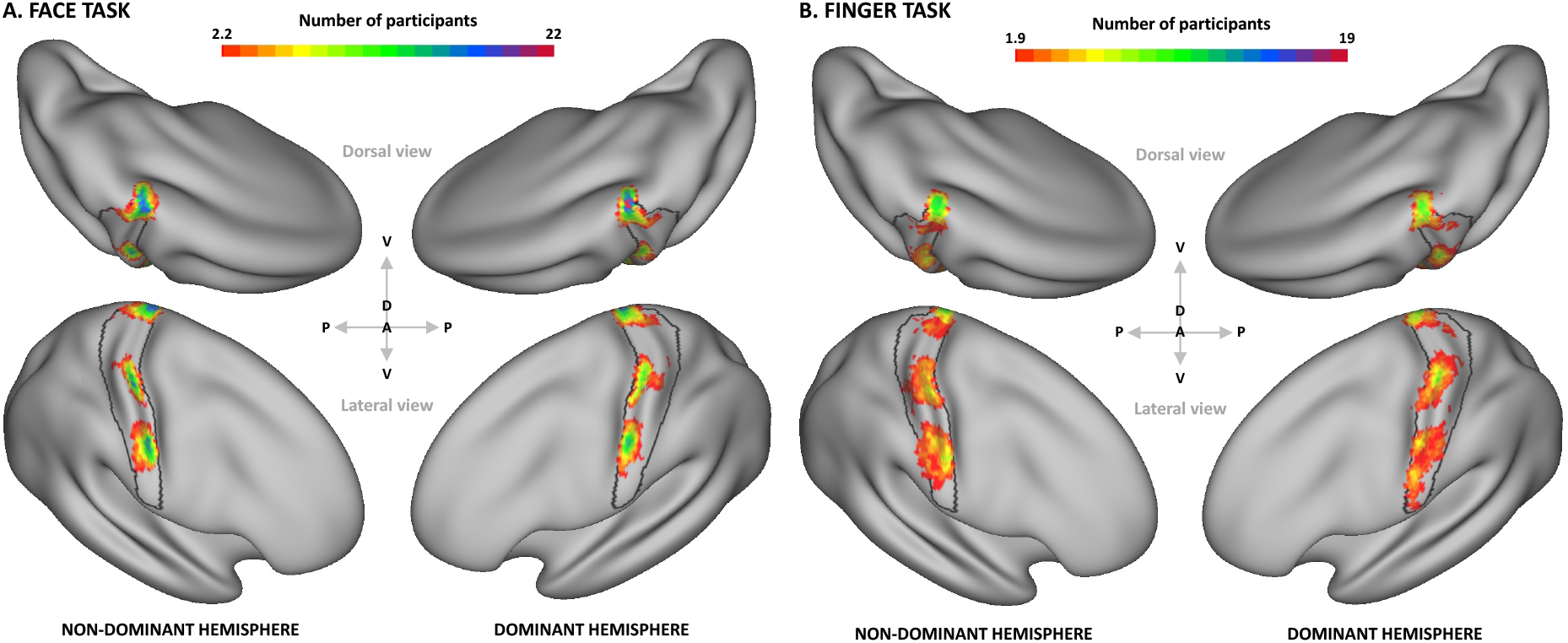
Consistency maps across participants of regions of interest within the primary somatosensory cortex (S1) used for the A) face and B) finger tasks respectively. Same legend as Fig. S2A.

### TWO ACTIONS DONE WITH THE SAME BODY PART CAN BE DECODED IN NON-PRIMARY AREAS OF THE HOMUNCULUS – non-dominant hemisphere

Activity levels evoked by different actions were significantly different in the primary ROIs of each body part (all *p* ≤ 0.013; Fig. S5A). Significant differences were also observed for arm movements in the Leg ROI (*t*_(21)_ = -6.04, *p* < 0.001) and for feet movements in the Hand ROI (*t*_(21)_ = -2.67, *p* = 0.014), and a trend was found for lip movements in the Leg ROI (*t*_(21)_ = 2.44, *p* = 0.024; other conditions: -0.27 < all *t*_(21)_ < 0.36, all *p* ≥ 0.722; Fig. S5A). Similar to the results observed in the dominant hemisphere, multivariate representational dissimilarities were significantly greater than zero not only in primary ROIs (all *p* values < 0.001, all *d* ≥ 0.99 CI [0.98 ∞]), but also in non-primary ROIs for arm and feet movements (all *p* ≤ 0.003, all *d* ≥ 0.66 CI [0.26 ∞]), and for lip movements in the Hand ROI (*z*_(21)_ = 196, *p* = 0.011, *d* = 0.55 CI [0.21 ∞]; Fig. S5B). These results suggest that action-related information content seems to be distributed across the Homunculus.

**Figure S5.**
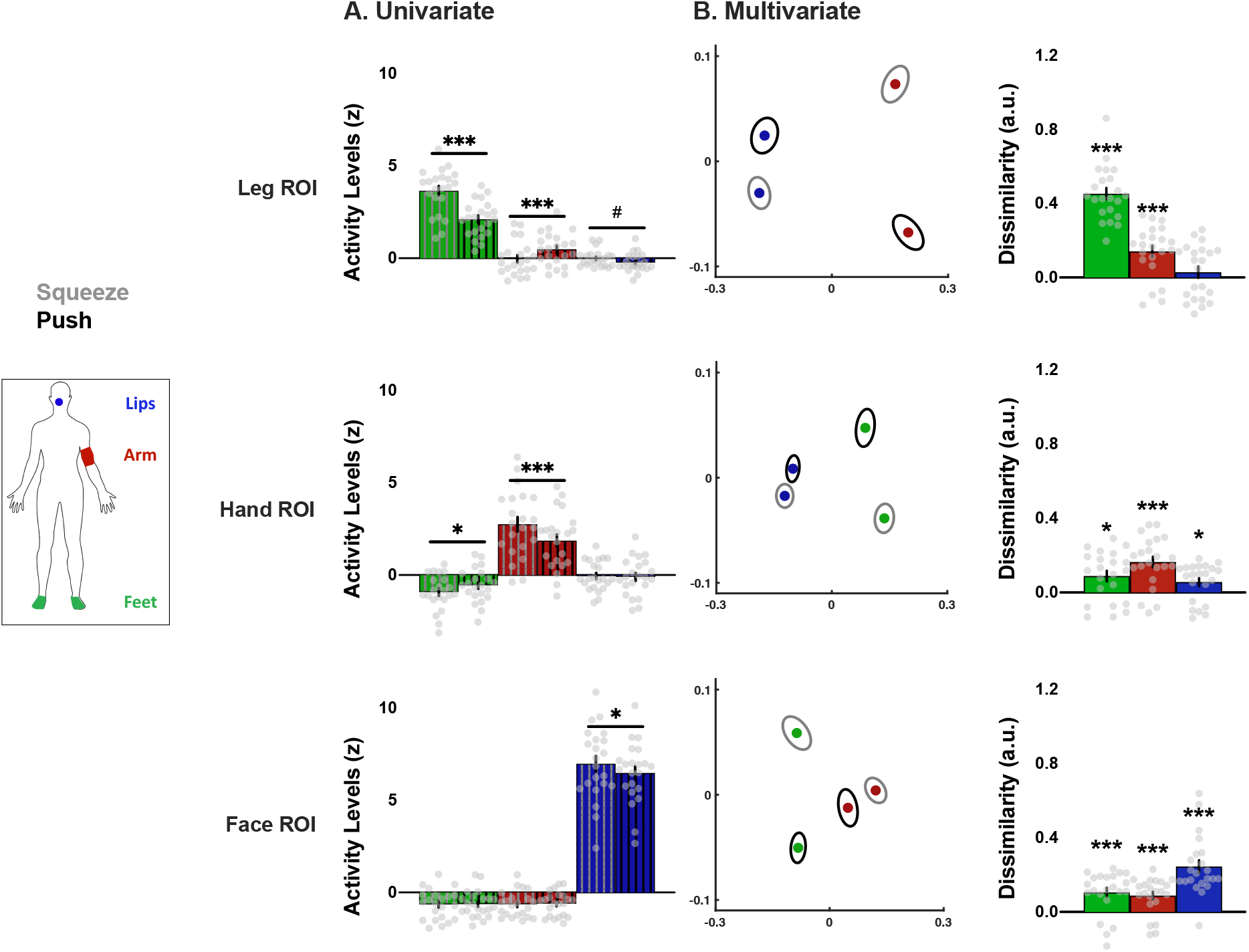
Univariate and multivariate content about different actions performed with a given body part across S1 Homunculus in the non-dominant hemisphere. All annotations are as denoted for Figure 4. \**p* < 0.017; *^#^p* < 0.033; \*\*\**p* < 0.001.

